# A Reference Feature based method for Quantification and Identification of LC-MS based untargeted metabolomics

**DOI:** 10.1101/2020.03.26.010769

**Authors:** Enhui Luan, Ken Cheng, Qiaoyun Long, Dehua Li, Zhenyu Li, Binghang Liu, Yalan Wang, Wei Li, Baosen Guo, Fengji Tan, Xiaoyi Yi, Lei Feng, Jiaping Song, Hancheng Zheng

**Author notes:** These authors contributed equally to this work (Enhui Luan, Ken Cheng).

## Abstract

Batch inconsistency is a major problem when applying LC-MS based untargeted metabolomics in real-time analysis situation such as clinical diagnosis or health monitoring. And inefficiency of collecting MS2 is a major problem for metabolite identification. Here, we developed a reference-feature based quantification and identification strategy (RFQI). In RFQI, samples are individually profiled using a pre-fixed reference feature table. Quantification results show that RFQI improves features’ overlap rate and reduce variance across batches significantly in real-time-analysis mode, and can find more than 4-fold numbers of features. Besides, RFQI collects MS2 from consecutive increasing samples for metabolite identification of pre-fixed features, thus it can effectively compensate for the poor efficiency of MS2 collection in data-dependent acquisition mode. In summary, RFQI can make full advantage of consecutive increasing samples in real-time analysis situation, both for quantification and identification.

## Introduction

Untargeted metabolomics is a wide-scale metabolic profiling technology which could broaden the possibility of novel metabolites discovery, and has been widely applied in the research of disease diagnostics, pathogenesis and disease prediction ^1,2^. Mass spectrometry coupled with liquid chromatography (LC-MS) is a commonly employed analytical approach, owing to its broad coverage of metabolites, high resolution and superior sensitivity^3^. However, as samples directly flow through the analytical platform, metabolic data collected from LC-MS varies between samples and batches in retention time (RT), chromatographic peak shape, ionization stability, mass accuracy (m/z) and signal responses, which could be caused by contaminants at inlet interface, ion suppression, differences in mobile phase and changes in performance of column, ionizer and mass analyzer ^4,5^. The instrumental variations become larger and more complex with increasing length of analytical experiments and number of batches.

To eliminate bias and increase comparability between samples and batches of untargeted metabolomic data, two methods based on XCMS are commonly used in research. One is to store samples until sample size reaches a proper number, then detect samples by LC-MS following XCMS analyzing all data together. This method is not suitable for real-time metabolomics analysis, for example, clinical or commercial health screening needs reports to be delivered to patients or customers immediately. The other is to analyze data from each batch independently and to align features of a batch with features of previous batches^3^. However, due to the loss of information in each process step for each batch, we have seen a low comparability of features and metabolites between batches, lots of missing values in the final feature table of samples and high coefficients of variation (CV) of feature intensities in QC **(Figure 4 & Figure S1)**.

Besides, the poor efficiency of metabolite identification based on MS2 is another bottleneck of LC-MS based untargeted metabolomics. It is mainly due to the limited number of known standards^6^ which is outside of our study aim, and the constraint number of collected MS2 with high quality, especially in data dependent acquisition (DDA) mode, in which only part of selected precursors is dissociated in one scan. Data-independent acquisition (DIA) mode can dissociate all precursors in one scan, but it increases difficulties in MS2 extraction of precursors and identification^6^.

Here, we developed a novel strategy RFQI (a Reference Feature-based method for metabolite Quantification and Identification) for untargeted metabolic data pre-processing, aiming to solve the poor-overlap problem of features across batches in real-time-analysis situation and improve the efficiency and accuracy in metabolite identification. Similar to the idea of assembling a reference sequence in genome and proteome, we firstly construct a reference feature table (rFT) which contains all possible features of certain bio-sample type. Multiple MS2 from incremental samples are then mapped to the rFT for feature annotation (**Figure 1**). Finally, the rFT was directly applied on each sample for profiling so that samples from all batches have a comparable feature list.

**Figure 1.**
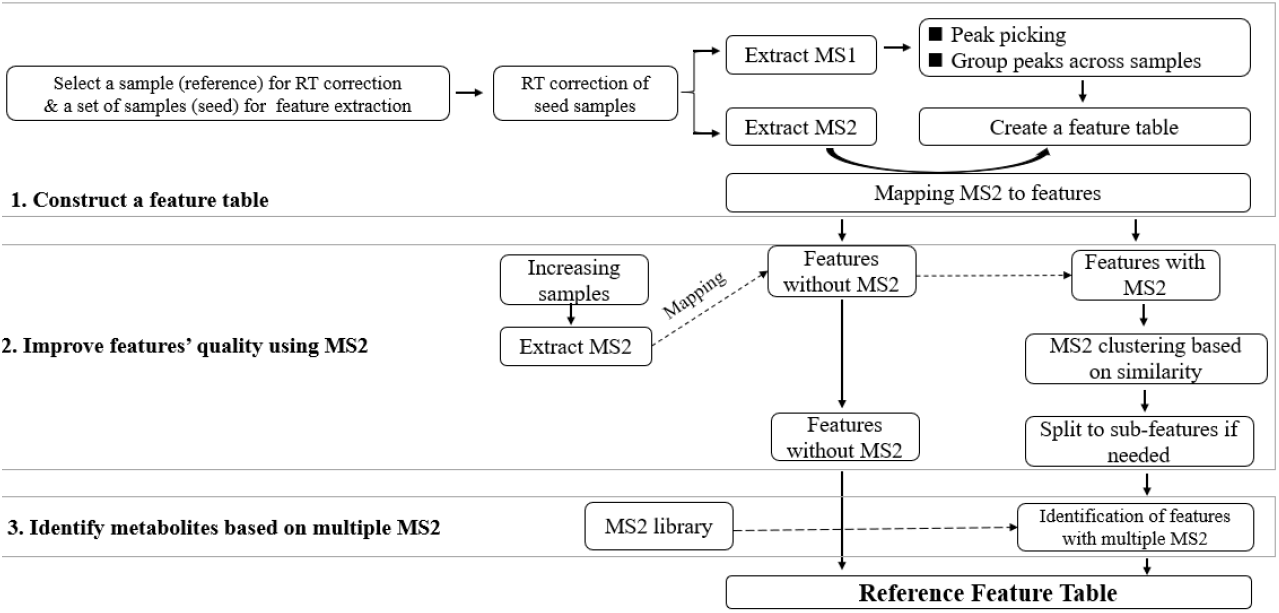
The working pipeline of RFQI to create a reference feature table (rFT).

Data from three consecutive batches including 148 samples originated from an untargeted metabolomics dataset (“ICXurineDB”) was used to evaluate the performance of RFQI, and 5831 urine samples measured across 140 batches in ICXurineDB were used to show the potential application of RFQI in real-time situation such as commercial. We found that the results of RFQI in feature detection, quantification and metabolite identification were largely improved. Additionally, while RFQI can satisfy the requirement of exporting analysis result in real-time, data mining results such as gender prediction based on increasing dataset has comparable even better performance than analyzing small batches altogether in research situation. In short, our findings show that RFQI could promote untargeted metabolomics to wilder application, such as clinical examination.

## Results

The term of “peak” refers to a mass spectral peak extracted by “Centwave”^7^ method in each sample, it represents a fragment of a precursor. The term of “feature” refers to a mass spectral peak with a unique m/z and RT which are obtained by assembling peaks from a certain number of samples^8^. Feature represents a precursor and could come from a metabolite protonated ion, adduct ion, fragment ion, dimers, trimers, isotope, instrument-specific ion or random noise ^3^.

### Introduction of RFQI strategy

RFQI is built based on three hypotheses:

1. The number of features in a specific type of sample, i.e., urine, is finite when applying a fixed detection method, since the number of both metabolites in bio-samples and their adduct types are relatively limited^9^. The whole feature set can be assembled by grouping feature information from a certain number of samples. We have seen that as the sample size increases, the feature number kept growing up and approached to a plateau (around 30,000 features) when sample number is over 75 (**Figure 2A**).
2. Collision induced dissociation (CID) of a precursor is not always stable due to the unstable CID energy, impurity, unstable detector, etc. **Figure S2** shows that the pattern of relative intensities of dissociated fragments derived from the same precursor are not stable. Additionally, many metabolites have similar precursors’ m/z and MS2 pattern, thus it may generate false discovery in metabolite identification when relying on only one MS2 spectrum.
3. We defined noise in LC-MS as randomly appearing signals rather than signals with low intensities, and the noise signals are only removed when the frequency of their occurrence in a large sample set are low (such as <50%). This may result in involving some background signals from mobile phase and instrument, but we think that these signals can be filtered during statistical analysis using variance threshold or missing value threshold, due to their relatively stability in each sample.

**Figure 2.**
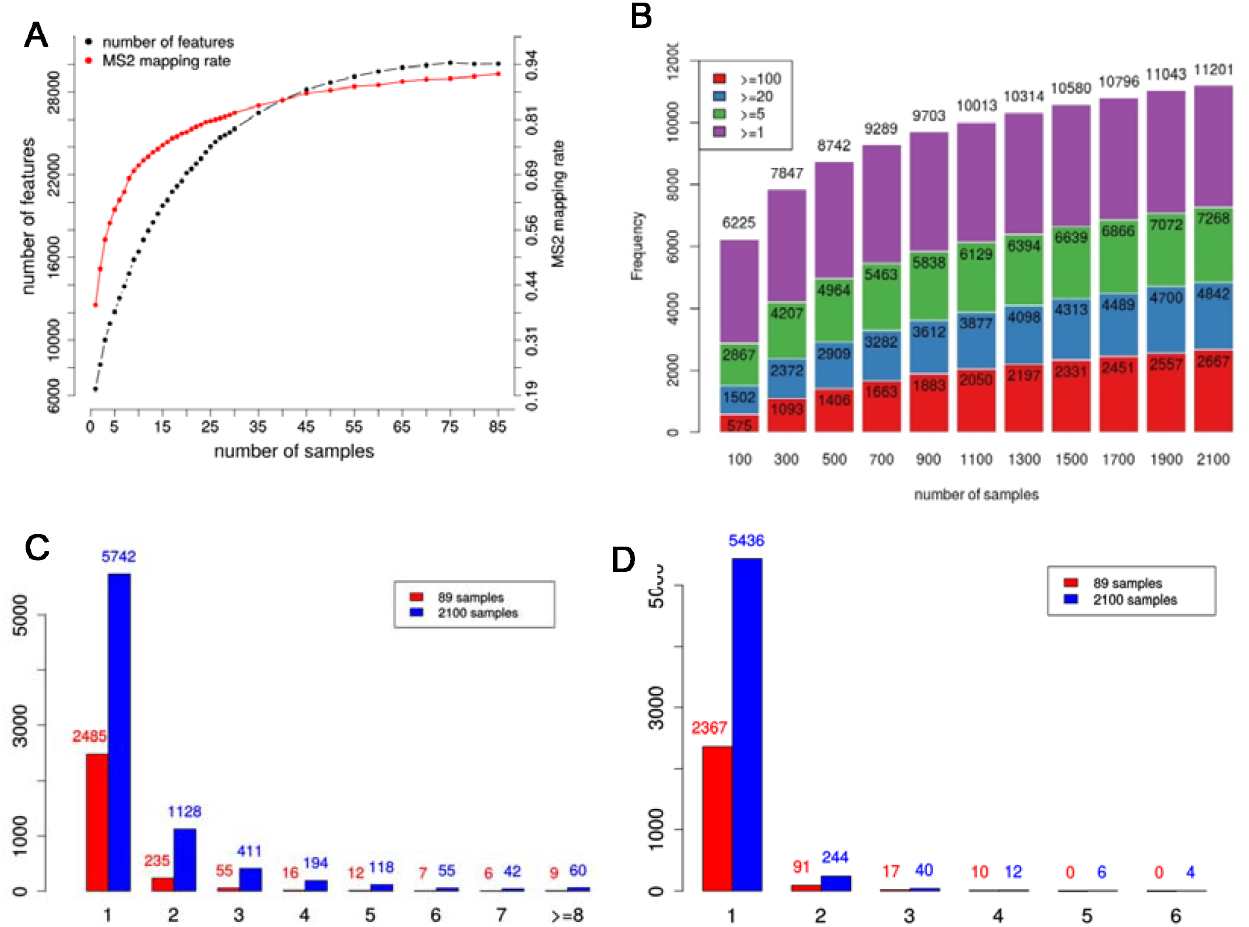
Validation of RFQI hypothesis. (A) The number of feature (black line) grows with the increase of sample number, and the proportion of MS2 mapping to features (red line) also grows with feature number increasing. (B) More features have mapped MS2 and features have more mapped MS2 when extracting MS2 from increasing samples. Colors represent different number of MS2 features having (1-5: purple, 5-20: green, 20-100: blue, and ≥100: red). (C) Number of features containing 1 to ≥8 MS2 clusters of MS2 when using MS2 from 89 samples and 2100 samples to map serial rFT. (D) Number of features which need to be combined between 1-6 features according to their MS2 similarities. The red bars show the results using MS2 from 89 samples, and the blue bar shows that using MS2 from 2100 samples.

Based on the above hypotheses, we firstly construct an rFT from a set of samples, then add metabolite annotation for this rFT using MS2 from a large amount of samples, finally we apply the annotated rFT to individual samples under the same condition (tissue type, sampling method, LC-MS/MS parameter) for profiling independent of batch. There are mainly three steps to build an rFT (**Figure 1**):

1. Construct a feature table from a set of samples A fixed sample (RT_reference sample) was selected for RT alignment and a set of samples (seed samples) were chosen for construction of rFT. We use Centwave method in XCMS to find peaks in each sample. To avoid introducing artificial bias in peak detection, we do not set strict threshold for peak picking in individual samples, such as noise level. A simple strategy is taken to group peaks into features, that peaks are treated as a rectangle in m/z-rt space. If peaks’ rectangles across different samples are overlapped or within pre-set threshold, they are defined as the same precursor and grouped as a feature. The m/z-rt area of a feature is the union of the original peaks rather than the intersection to cover the shift in RT as much as possible.
2. Check and improve the features’ quality using multiple MS2 MS2 from samples are mapped to the above feature table using precursors’ m/z and rt. With the increasing number of samples, more features will have MS2 and the number of MS2 belonging to one feature will also increase. For example, MS2 from 148 samples were mapped to 4076 features in rFT, in which 3046 features had more than two MS2, while MS2 from 2100 samples were mapped to 10176 features, in which 7011 features had more than two MS2 (**Figure 2B**). The fact that one feature can have many MS2 facilitates us to check the features’ purity by MS2 similarity clustering. Similarity of MS2 belonging to the same feature and among different features were checked based on MS2 from 148 and 2100 samples, respectively. We found that the majority of features with more than three MS2 (88.0% and 74.1% when using MS2 from 148 and 2100 samples, respectively) had consistent MS2 (**Figure 2C**) which indicates these features were derived from one precursor. While the similarity of MS2 belonging to different features was rather low, only 4.7% (using MS2 from 148 samples) or 5.3% features (using MS2 from 2100 samples) were similar in MS2 spectrum and may need to be combined (**Figure 2D**)
3. Identify features based on multiple MS2. All MS2 mapped to features are applied for metabolite identification. There are two advantages of using multiple MS2 but not selecting one representative MS2 for identification. On one hand, this can cover the variation of CID as much as possible and increase the probability of matching to the reference MS2 from MS2 library such as NIST. On the other hand, when comparing with library MS2, features with multiple MS2 will generate a vector of similarity scores rather than a scalar, so we can identify metabolite more accurately by comparing similarity score distribution. The algorithm in detail for metabolite identification can be found in method part.

### Features extracted using RFQI had better repeatability

Two rFT were constructed by using 85 samples from batch1 (serial rFT) and 85 samples randomly selected from three batches (random rFT), respectively. We applied both of the two rFT to profile the samples from the three batches.

There were 30018 features in the serial rFT. Among them, 29544, 28700 and 29043 features were detected in >50% samples in Batch 1-3, and 28430 features (94.7%) presented in >50% samples of all three batches (**Figure 3A-C, blue rectangles**). By comparision, features in three batches were seperately extracted using XCMS and 7036, 7776, 6759 features were presented in >50% samples in each batch **(Figure 3A-C, red rectangles)**. Among them, only 3866 features (~53.8%) were shown up in all three batches **(Figure 4D)**. These results show that RFQI can use more information of the raw data and the repeatability of features detected across batches were significantly improved by RFQI.

**Figure 3.**
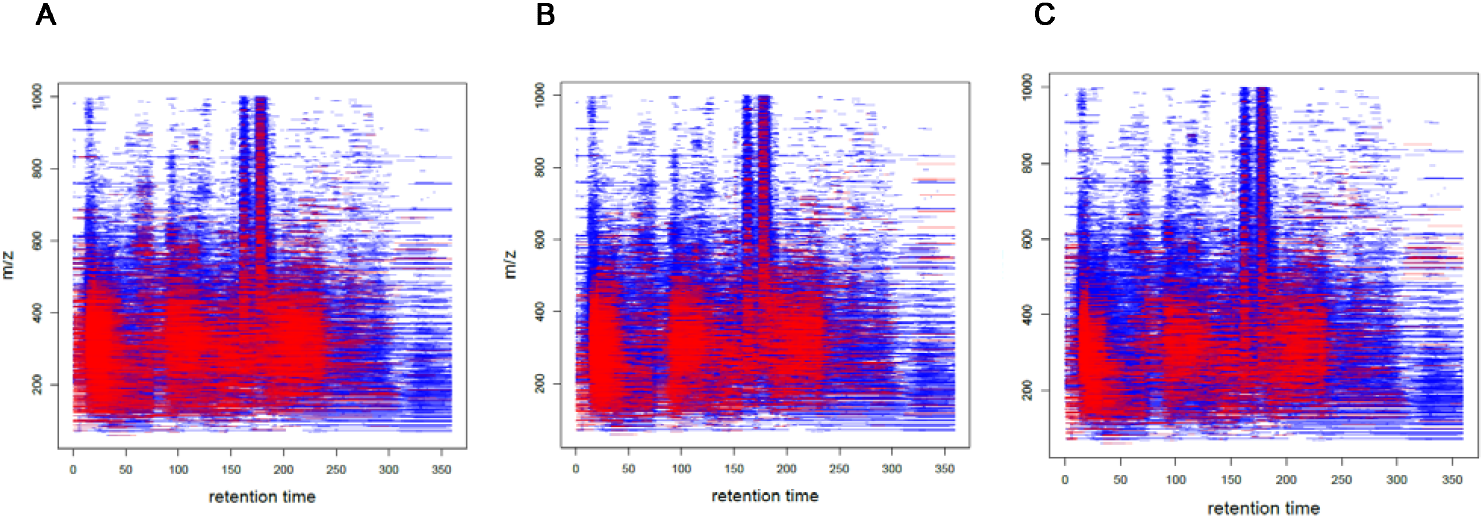
RFQI can use more information. Features extracted by RFQI (blue rectangles) and XCMS (red rectangles) using the data from the 3 batches, respectively (A-C).

To examine the hypothesis 1 mentioned above, we compared the random rFT with the serial rFT. Most of the features are overlapped **(Figure S3A)**. Random rFT had 29519 features, which is close to serial rFT’s 30018 features. One-to-one overlapped features (features in serial FT only map to one feature in random rFT, and vice versa) were extracted to compare the consistency in sample quantification, we found that most of the Pearson Correlation Coefficient were greater than 0.9 **(Figure S3C)**. Both rFT had their unique features **(Figure S3B)**, whereas these unique features have relatively low intensity **(Figure S3F)**, more missing values **(Figure S3D)**, and higher QC’s CV **(Figure S3E).**

### Features extracted using RFQI had better stability in Quantification

Profiling results of three batches using serial rFT were used to check the feature stability in quantification. Firstly, we checked the stability of feature quantification by calculating the CV of features intensity in QCs within one batch. 70.6%, 71.4% and 70.2% features having CV<30% extracted using RFQI in Batch 1-3 (median CV: 19%, 17%, 18%) respectively, while 53.7%, 61.5% and 60.3% features having CV<30% using XCMS (median CV: 28%, 25%, 25%)(**Figure 4A-C**). The CV of features’ intensity in QCs was much smaller using RFQI than using XCMS (P-values: 2.2×10^−71^, 9.4×10^−11^, 4.8×10^−7^).

**Figure 4.**
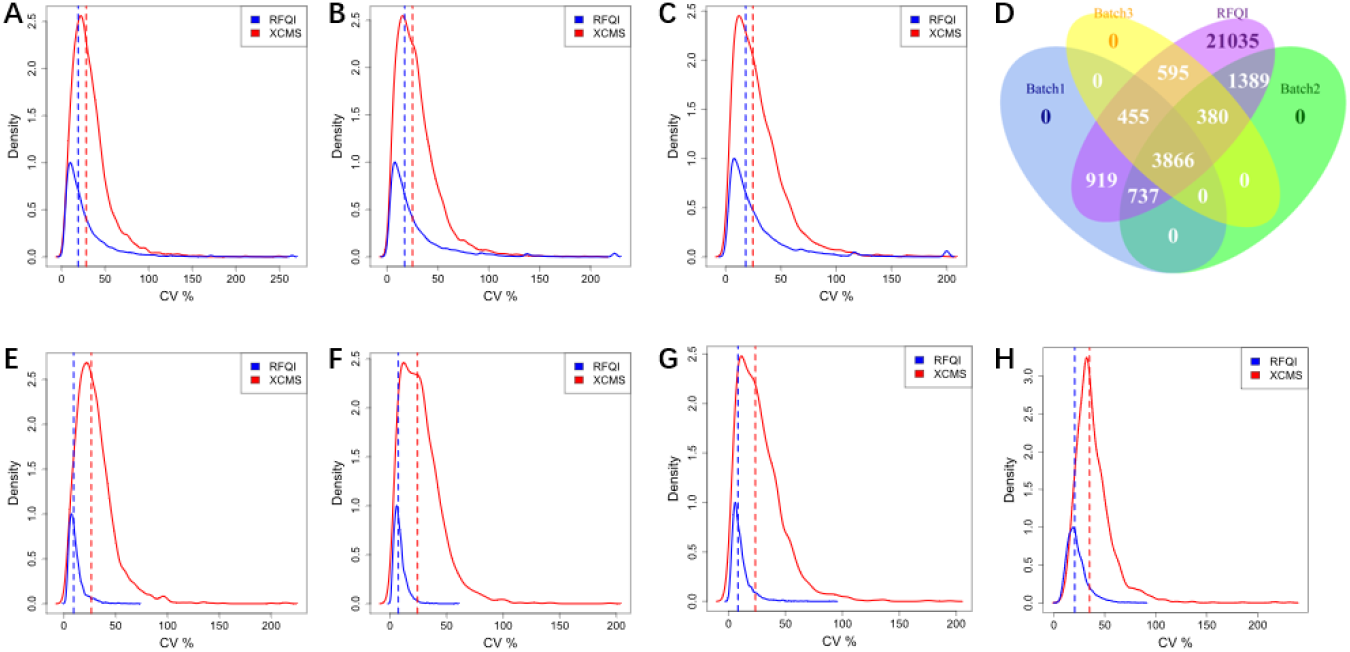
Comparison between RFQI and XCMS in feature stability within and between three batches. (A-C) Coefficient of variance (CV) distribution of feature intensity in QCs from Batch 1-3, respectively. D. Feature area overlap of RFQI and XCMS for three batches (XCMS features from each batch are first aligned with RFQI’s, then using RFQI-mapped features to do feature comparison between batches. One feature in RFQI may correspond to more than one feature in XCMS, and vice versa.). (E-G) CV distribution of the one-to-one overlapped features in RFQI and XCMS (n=2088) in QCs from Batch 1-3, respectively. (H) CV distribution of overlapped features (n=2088) in QCs from all three batches (blue lines: RFQI, and red lines: XCMS).

Since the number of features extracted by RFQI and XCMS were largely different, the stability of one-to-one overlapped features (n=2088) in RFQI and XCMS were further compared. We have seen that 96.7%, 99.3%, 96.7% features having CV<30% using RFQI in Batch 1-3 (median CV: 10%, 7%, 8%), respectvley, while 58.3%, 63.9% and 63.3% features having CV<30% using XCMS in Batch 1-3 (median CV: 27%, 25%, 25%), respectively (P-values: 1.1×10^−281^, 1.7×10^−298^, 4.3×10^−217^; **Figure 4E-G**). The differences in CV of feature intensity were more obvious when comparing the overlapped features (n=2088) in QCs from all three batches (**Figure 4H**), 82.9% and 32.0% of features having CV<30% in RFQI and XCMS (median CV: 20% and 35% in RFQI and XCMS, P-value=1.6×10^−229^), respectively. Our findings suggest that RFQI had improved performance in feature quantification within and between batches.

### More metabolites were identified using RFQI

Traditional method for metabolite identification is to select one representative MS2, such as MS2 corresponding to the maximum precursor’s intensity, and assign a hard threshold to filter possible false discovery^6,10^. In RFQI, all MS2 derived from a large number of samples and mapped to one feature are applied for this feature’s metabolite identification, which can increase the possibilities of mapping them to library MS2. Besides, as RFQI groups peaks into features by union rather than intersection, the area of a feature is wider so that more MS2 can be mapped to this feature.

For traditional method, we assigned similarity score > 0.9 as threshold. And for RFQI, as there are many similarity scores generated by comparing multiple experimental MS2 with one library MS2, we defined a feature is identified when there is at least one similarity score greater than 0.9.

138 metabolites were identified by RFQI using MS2 from the 148 samples in three batches. When using traditional method, there were 89, 82, and 80 metabolites identified in Batch 1-3, respectively and only 40 metabolites were detected in all three batches, which were also identified in RFQI result **(Figure 5A)**. Since there were more features having MS2 in the rFT compared with XCMS result, we further compared the identification result of one-to-one mapping features (n=2088) that detected in both three batches by RFQI and XCMS. RFQI identified 44 metabolites, while traditional method identified 29 **(Figure 5B)**.

**Figure 5.**
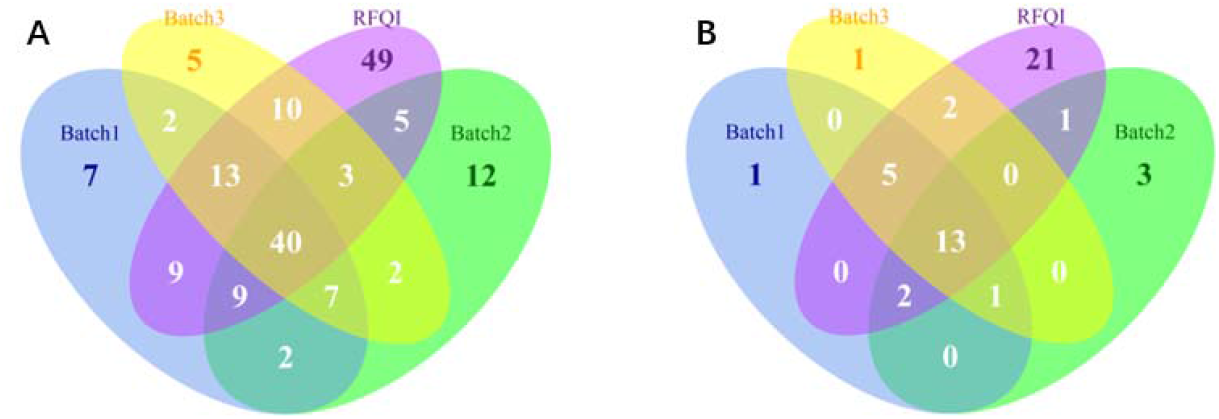
RFQI identifies more metabolite. A.Venn-diagram of metabolite numbers identified in Batch 1-3 using traditional method (Batch 1-3) and all three batches using RFQI (RFQI). B. Venn-diagram of metabolites numbers identified using traditional method (Batch 1-3) and RFQI (RFQI), using the one-to-one mapped features in RFQI and XCMS (n=2088).

MS2 information of features in rFT can be further enlarged with the increase of sample size (**Figure 2B**), thus, metabolite identification in rFT could be continuously improved in future experiments. Compared to 138 metabolites identified using MS2 from 148 samples, when taking MS2 from 2100 samples, the number of identified metabolites increased to 430 in urine sample. This is another strong merit of fixing an rFT of a specific sample type which improves the possibility of certain feature having mapped MS2 from a larger sample set, which can effectively solve the limits of DDA mode, and more importantly enhance the performance of metabolite annotation. However, we found that as the sample size reached to 2100, the number of features having MS2 stopped increasing. Thus, we propose that for the features without MS2 in rFT, their MS2 can be targeted collected in the future experiments, to increase the power for metabolite identification.

### RFQI had a higher sensitivity in metabolite identification

As stated above, for one feature with multiple MS2 mapped, RFQI generates a vector of similarity score by comparing multiple experimental MS2 with one library MS2 rather than a scalar. This offers a strategy to control false discovery rate based on hypothesis testing for two vector distribution rather than setting a hard threshold such as >0.9.

For example, feature FT01143 in the serial rFT had 162 MS2, so for each MS2 we can get a vector of similarity scores with 161 elements by comparing this MS2 with the rest of MS2 belonging to the same feature (red lines in **Figure 6C**). The distribution of a vector can represent the probability of occurrence of the precursor generating this MS2 and if the mean or median of vector A (represent A MS2) is significantly greater than that of vector B (represent B MS2), we define the probability of A MS2 belonging to the feature is greater than B MS2. Under this rule, although MS2 of L-Pyroglutamic acid (L-Py) from NIST was similar to FT01143 MS2 when using a hard threshold (average similarity score > 0.99), only less than 5% MS2 belonging to FT01143 (**red lines in Figure 6C**) were smaller than L-Py (**green line in Figure 6C**), so we concluded that the probability of L-Py being the feature less than 0.05 and thus, FT01143 was not identified as L-Py (**Figure 6C**).

**Figure 6.**
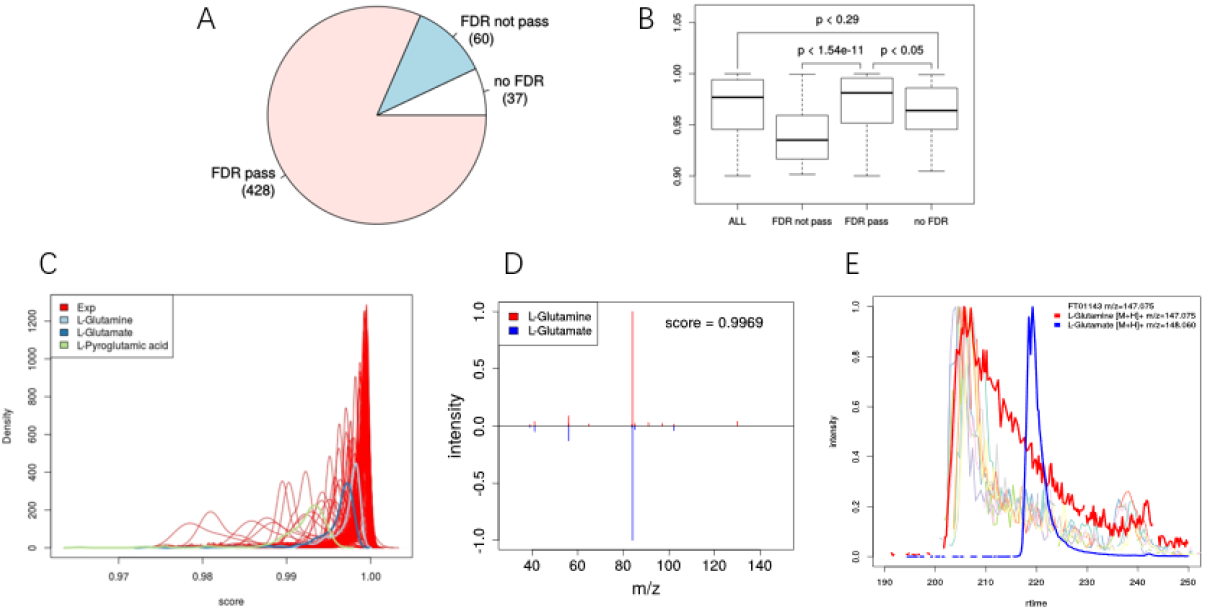
RFQI offers a highly sensitive method for metabolite identification. A. Filtering metabolites identified by hard threshold of similarity score >0.9 using RFQI. “No FDR (37)” means that 37 features have less than three MS2 (<3). “FDR pass” means that RFQI recognizes the feature identification as true positive result. “FDR not pass” means that RFQI defines the feature identification as false positive result. B. Max similarity score distribution of FDR filtering result. C. similarity score distribution of all experimental MS2 belonging to the same feature (FT01143) (red lines; each red line represents a similarity distribution of one experimental MS2 compared with the rest of experimental MS2) and similarity score distribution of all experimental MS2 with library MS2 (blue line: L-glutamate; light blue line: L-glutamine; green line: pyroglutamic acid). D. MS2 patterns of L-glutamate (blue) and L-glutamine (red) which are similar. E. The extracted ion chromatograph (EIC) of L-glutamate (blue line) and L-glutamine (red line) standards, as well as the peaks of FT01143 in many samples (other lines).

Besides offering a strict strategy for reducing false discovery, RFQI can even distinguish metabolites that have very similar MS2 and precursor m/z. this is often the case of metabolites, because structural similarity is a universal paradigm for small molecules. For example, L-glutamine (Gln) and L-glutamic acid (Glu) have similar molecular ion weight ([Gln+H]+ and [Glu]+), and MS2 patterns (**Figure 6D**), except slightly difference in m/z intensities in MS2 (**Figure 6D**). RFQI identified FT01143 as Gln because the probability of Gln was greater than Glu (**Figure 6C**). To validate the result, pure Gln and Glu standards were analyzed by LC-MS/MS, the result showed that L-Glutamine was more similar to FT01143 both in RT, molecular ion weight, and distribution of isotopic ion peaks (**Figure 6E**).

We further examined the false discovery rate of hard threshold 0.9 by using MS2 from 2100 samples. We defined an identification as false discovery if the probability is less than 0.1. About 525 features had MS2 whose max similarity score with library MS2 is greater than 0.9. Among them, only a small set of features had less than three MS2. While most features passed RFQI checking, 60 features were failed (**Figure 6A**).

RFQI offers a soft threshold for identification. As showed in **Figure 6C**, even though similarity score distribution of “FDR pass” and “FDR not pass” features are significantly different, some features with low similarity score are conserved, and some features with high similarity score are discarded. As for features that cannot be checked due to lack of more MS2, we suppose them having the same FDR as features with more MS2 because the distribution of similarity score of both (no FDR and overall in **Figure 6B**) are similar.

## Discussion

Real-time-analysis situation such as clinical requires reports delivered in time once samples are received. So it is necessary to analyze metabolomics data batch by batch even sample by sample while keep them comparable. At present, targeted absolute quantification of biochemicals is still the mainstream in clinical medical test, however, this method only uses a little information of bio-sample. LC-MS based untargeted metabolomics is a powerful tool in grasping information of large-scale metabolome in biological system and has been widely applied in scientific research. However, when analyzing data batch by batch across a long duration, due to signals consistency of peak shape, m/z and RT between samples and batches, it is difficult to align samples by features.

To solve this problem, we developed RFQI, a new strategy of metabolic profiling by individual sample based on pre-fixed rFT of a specific sample type. This approach can effectively avoid pre-processing samples from batches altogether. Our results demonstrate that features extracted from multiple batches using RFQI have improved performance in repeatability and stability in quantification.

In addition, fixing a reference feature table offers an additional benefit, i.e. we can collect MS2 from hundreds even thousands of samples for a feature. On one hand, this can make more features having MS2, which will be helpful for feature annotation. On the other hand, one feature can have multiple MS2, this offers a distribution-based identification method which can largely improve the accuracy of identification, even for distinguishing metabolites with similar structures.

We applied RFQI on a part of urine untargeted metabolomics dataset from ICXurineDB (n=5831) for metabolic profiling, based on a reference feature table created using 90 samples randomly selected from the same dataset. The results in quantification can be used to predict phenotypes of participants, such as gender **(Figure S4)** effectively. Our findings are in accordance with the others^11^, which further demonstrates that RFQI is an effective tool for untargeted metabolomics data processing and could be potentially applied in clinical situation for following data mining.

Nevertheless, there are still some problems in untargeted metabolomics studies for both clinical situation and research community. For example, data normalization across batches is usually relying on QC’s stability (CV) ^12^, but this only can demonstrate that normalization method minimize variation, not only for QC sample, but may also for biological samples that may have difference. Splitting the QC dataset into training-set and testing-set can effectively avoid over-normalization, but it is inevitable to make a standard dataset with given concentration differences. Moreover, although RFQI can collect more MS2 for more features from a large number of samples, it is still difficult to identify all features due to lack of standards.

In conclusion, RFQI is a total novel strategy for untargeted metabolomics data pre-processing which can enlarge the potential medical applications of untargeted metabolomics in future.

## Methods online

### Urine sample dataset

A urine dataset obtained from ICarbonX database (ICXurineDB), Shenzhen, China was applied in the study to describe and verify the RFQI algorithm. The database contains 5831 urine untargeted metabolomics data from Meum products until December 2018 and additional 1200 QC samples, which were separately analyzed in 140 batches over about 1.5 years. The study was approved by the Institutional Review Board (IRB) at ICarbonX (IRB2019003, Shenzhen, China) and participants have signed informed consent before sampling.

To show the performance of RFQI in data pre-processing, data from three continuous batches (Batch 1 contains 7 QCs and 82 samples, Batch 2 contains 5 QCs and 31 sample, and Batch 3 contains 4 QCs and 19 samples, respectively) were randomly extracted. To verify the application of RFQI in a large-scale dataset, data from all 140 batches were taken.

### Sample collection and LC-MS based untargeted metabolomics analysis

Urine sample were collected by participants themselves using a urine sample collection kit (Meum, IcarbonX, Shenzhen, China). An approximate 5 mL urine sample in tube were transported at room temperature within 24 hours and frozen immediately at −80°C when arriving at laboratory until analysis.

Urine samples (100 μL) were diluted with water (300 μL), vortex for 30 s, and ultra-sonicated for 10 min at 4°C. After centrifugation at 15000 *g* for 15 min at 4°C, the supernatants (100 μL) were transferred to LC vials for sample analysis. A pooled urine sample of 300 healthy people was prepared as a quality control sample (QC). QC were injected regularly (5 injections at the beginning for equilibration, at the end and at every 8^th^ injection in each batch sequence) throughout all batches to monitor the instrumental stability.

LC-MS based untargeted metabolomics was conducted in ultra-high-performance liquid chromatography (UPLC) system (ExionLC-AD, Sciex, Concord, New Hampshire) connected to a quadropole-time-of-flight mass spectrometer (TripleTOF 5600^+^, Sciex) via an electrospray ion source (ESI) interface. Sample solution (2 μL) were injected into a HILIC column (Acquity UPLC BEH Amide 1.7 μm, 2.1×50 mm, Waters, Milford, MA) operating at 30°C. A binary eluent system (eluent A: 25 mM ammonium acetate and 25 mM ammonium hydroxide in water; eluent B: 100% ACN) was applied at 0.6 mL/min of flow rate. The gradient profile was 0 – 4 min: 5% to 35% A, 4 – 4.5 min: 35% A, 4.5 – 4.6 min: 35% to 5% A, and 4.6 - 6 min: 5% A. The ESI was operated at 5000 V in positive mode using nitrogen as nebulizing and drying gas. The temperature of gas at ESI source was 600°C and the gas pressure was 60 psi. The de-clustering potential (DP) was 100 V. Data were acquired in data dependent acquisition (DDA) mode. Performing a full scan, mass spectra were recorded at 50 - 1000 m/z for 150 ms, using a collision voltage 10 V. Performing a DDA MS/MS scan of the five most abundant mass peaks in the full scan, mass spectra were recorded at 25 – 1000 m/z for 50 ms per acquisition, using a collision voltage 30 eV. Data were collected in profile mode by using Analyst TF 1.7.1 (Sciex) software, and converted to mzXML using MSConvert (PreteoWizard 3.0.18285, downloaded in Jan 2017).

ACN, MeOH and water (Fisher, USA) in HPLC-MS grade were used in the study. Ammonium acetate and ammonium hydroxide were purchased from Merck (Germany).

### Metabolite library

The metabolite library using for metabolite identification was laboratory-owned database which was established by injecting standard compounds to the same model of instrument (Sciex 5600+) with the same parameter setting and web-available database, such as the Human Metabolome dataset (HMDB), METLIN, ChemSpider, SciFinder and NIST. At present, the database contains MS/MS spectra of 1471 metabolites.

### Procedure of RFQI

The procedure contains (1) constructing a reference feature table (rFT) (2) checking and improving the rFT quality using multiple MS2 (3) identification of features based on multiple MS2 (**Figure 1**); (4) metabolic profiling of samples from batches based on the rFT.

#### Constructing a reference feature table

We firstly selected one QC sample from ICXurineDB with good performance in chromatography as a reference for RT correction (“ref_RT”), and then selected a set of samples as seed samples. After RT correction of seed samples aligning with ref_RT sample using “obiwarp” algorithm (binSize=0.25)^8^, peaks in seed samples were individually extracted using “Centwave” algorithm (ppm=10, peakwidth=c(2,30), snthresh=3, noise=0) in XCMS. To group peaks into features across the seed samples, we treat peaks as a rectangle defined by [mzmin,mzmax,rtmin,rtmax]. If two peaks have overlapped or adjacent mz range and rt range, they are grouped together. The grouping step was done iteratively. The final feature area is the union of peaks belonging to it to cover the potential shifts as much as possible. This grouping algorithm is different from groupChrompeaks_nearest and groupChrompeaks_density which take median RT and m/z of peaks for grouping.

#### Extracting and Mapping MS2 to reference feature table

Samples’ retention time were firstly aligned with the above ref_FT sample using “obiwarp” algorithm in XCMS. MS2 spectrum were then extracted from samples and mapped to reference feature table. An MS2 is mapped to a feature only when it’s precursors’ mz and rt fall into the area of a feature.

#### Checking and improving rFT quality using multiple MS2

Features in rFT could have multiple MS2 and with the increasing of sample size, more feature will have MS2 and the number of MS2 belonging to one feature also increases. According to the number of MS2, features can be divided into three types: features with >=3 MS2, features with <3 MS2 and features without MS2. For the features with more than three MS2, MS2 similarity clustering^13^ can be calculated to evaluate and improve the features quality. Cosine distance is used to measure the similarity of paired MS2 and ‘dbscan’ method to cluster MS2 according to similarity scores. We define two MS2 similar if their cosine similarity score >0.9 and the minimum neighbor is 3. If the multiple MS2 spectra belonging to one feature can be divided into more than one cluster, the feature will be split into sub-features.

**Figure 7** shows two conditions of features by checking their MS2 similarity: a pure feature (**Fig. 7A-C**), an impure feature which can be clearly split into two clusters (**Figure 7D-F**). In the data from Batch 1, 3231 out of 30058 features contained MS2 information using MS2 from 85 samples, while 2991 out of the 3231 features with MS2 were pure features and 240 were split into 2 sub-features.

**Figure 7.**
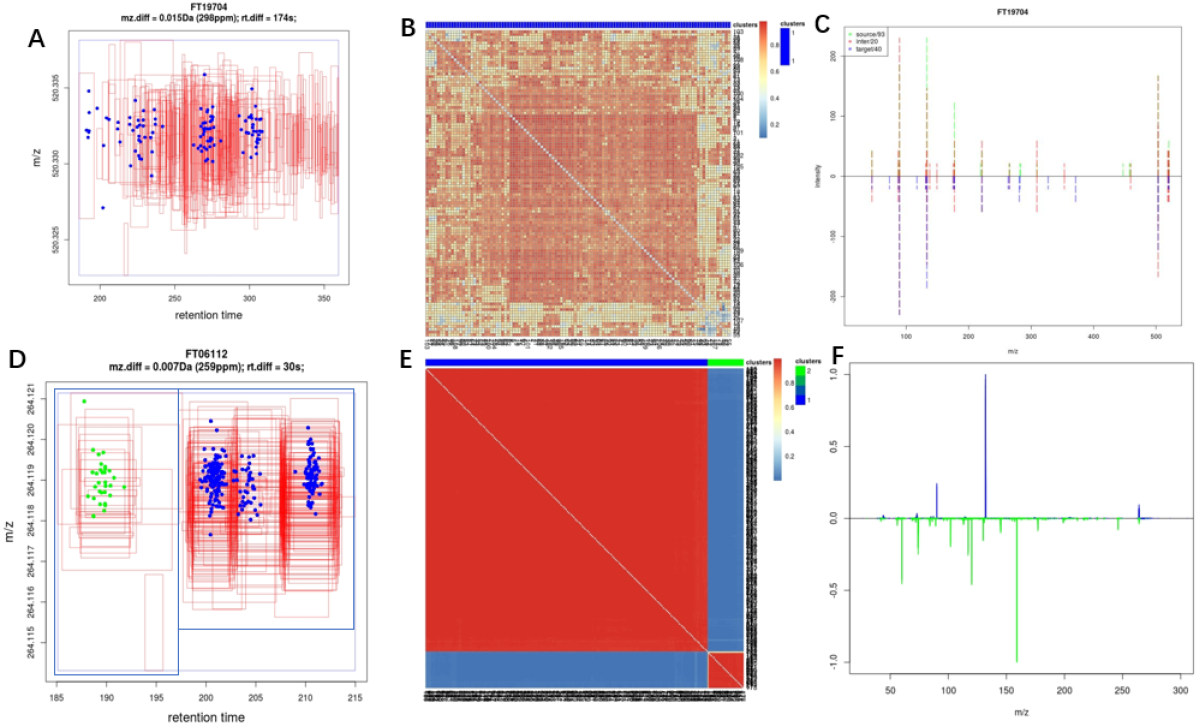
Check the purity of features by using multiple MS2. <u>Figures A, B, C – a pure feature (RT span 174s, m/z 298) as an example.</u> (A) RT-m/z plot. The red frames refer to the features in all ref_samples, and the blue points refer to the mapped MS2. (B) heatmap * of all MS2 belonging to the feature. (C) comparison of MS2 from the same precursor. <u>Figures D, E, F – a feature (RT span 30s, m/z 259) having two clusters of MS2.</u> (D) RT-m/z plot. The red frames refer to the features in all ref_samples, and the blue and green points refer to the mapped MS2 in clusters. (E) heatmap of all MS2 belonging to the feature. (F) comparison of MS2 from different clusters in D and E. * The score of heatmaps in the figure were calculated by cosine similarity, and clusters are defined by dbscan

For features without MS2 and less than three MS2, MS2 information can be targeted collected in new experiments to enlarge the information of rFT.

#### Identification of metabolites based on multiple MS2

The essential principle of metabolite identification is to compare the similarity between experiment-sourced MS2 (Exp_MS2) and library-sourced MS2 (Lib_MS2). In RFQI, multiple Exp_MS2 belonging to the same feature were applied for metabolite identification.

The procedure of identification was shown as below

1. For one feature with M (M >2) Exp_MS2, we compute the cosine similarity score of each MS2 with other MS2, so we can get M vectors whose length is M-1 **(Figure 8 left**).
2. For one feature, if m/z of a certain type of adduct of one metabolite falls into the feature’s m/z range, we select this Lib_MS2 of this metabolite to compare similarity with all Exp_MS2 belonging to this feature. One feature may have N mapped metabolite, for each mapped metabolite, we get a vector of similarity score of length M **(Figure 8 right)**.
3. For each mapped metabolite, we compare the Lib-MS2&Exp-MS2 similarity vector with each Exp-MS2&Exp-MS2 similarity vector using t test, if the mean value of Lib-MS2&Exp-MS2 vector is not significantly less than one Exp-MS2&Exp-MS2 vector, we define this Exp-MS2 as a pro that favours with the feature being the metabolite. So for each metabolite mapping to a feature with M MS2, we can get x pros and y cons, where x plus y equals M. x/M means the probability of the metabolite’s Lib-MS2 belonging to the feature. If x/M is greater than a threshold (such as the commonly used α=0.05), we conclude the feature is derived from the metabolite.
4. If many metabolites mapped to one feature, we can compare each x/M and choose the one with higher probability.
5. For features with less MS2 (<3), we compare each of Exp-MS2 with mapped Lib-MS2 and take the maximum similarity score. We use a hard similarity score threshold for this situation, but as samples increasing, this feature can have more MS2 and use the strategy above.

**Figure 8.**
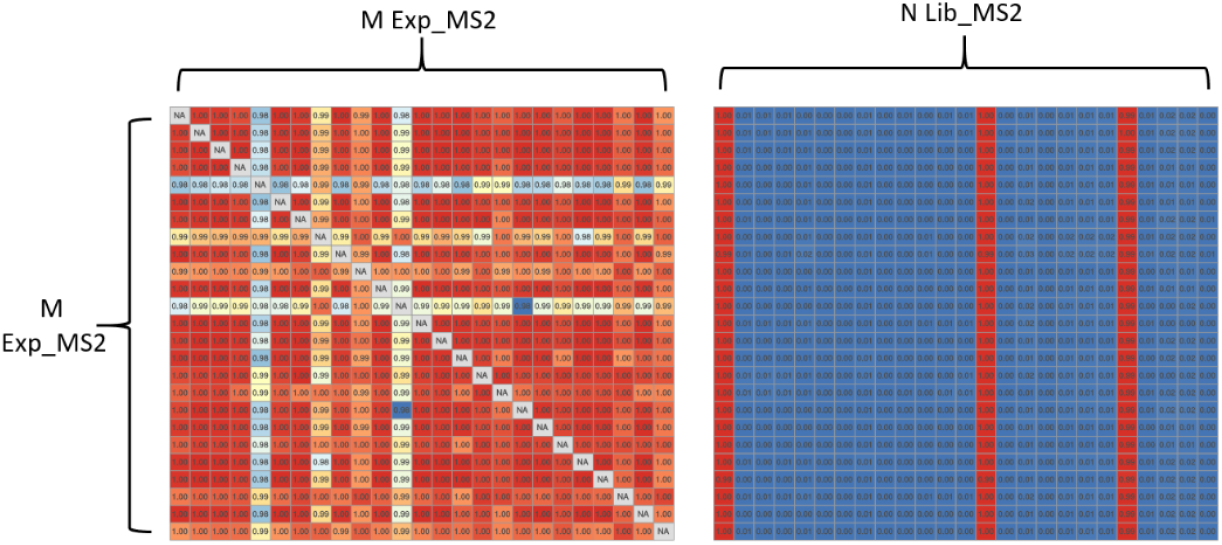
Principle of feature Identification by using multiple MS2. Left: a matrix of similarity score between Exp_MS2 and Exp_MS2 (n= M). Right: Lib_MS2 with Exp_MS2 similarity score matrix. M represents that a feature has M MS2 and N represents MS2 of N metabolites were mapped to the feature based on m/z. Each column is a similarity score vector. And each column in the right matrix compares with all columns in left matrix using t test.

#### Metabolic profiling for samples from batches based on the rFT

Metabolic features in samples from batches can be individually profiled based on the ranges of mz and RT in the ref_FeatureTable, mainly following the steps: (1) RT correction using the ref_RT as reference and “obiwarp” algorithm (binsize=0.6, centerSample=ref_RT); (2) integration of features at the defined m/z and RT ranges in rFT. To be noticed, the samples for profiling should be in the same type as that in rFT.

### Other approach for data pre-processing

Data from Batch 1-3 were separately pre-processed using the XCMS R package (version 3.6.0)^14^. The parameter settings for “Obiwarp” algorithm (binsize=0.25, centerSample=ref_RT) and “Centwave” algorithm (ppm=10, peakwidth=c (2,30), snthresh=3, noise=0) were the same as RFQI, while that of “groupChromPeaks-density” algorithm (minFraction=0.5, bw=30, mzwid=0.25) for grouping and “FillChromPeaksParam” algorithm (default settings) were set under manual optimization.

### Other approach for metabolite identification

A representative Exp_MS2 spectrum of a feature was selected for metabolite identification. Here we use the “max intensity” strategy, that the selected MS2 has the largest precursor intensity. The cosine similarity between Exp_MS2 and Lib_MS2 was then calculated as above.

### Classification of gender

Samples (n=5831) from ICXurineDB were used to check the potential application of RFQI in large-scale dataset. The distribution of gender is balanced, including 3021 male and 2810 female. There are more than 30,000 features profiled using RFQI, and 10649 features which have multiple MS2 were used for gender classification. We randomly selected 70% of data as training dataset and 30% as testing dataset. Feature selection is based on training dataset using student t-test (P-values were adjusted by “p.adjust” function in R with “fdr” method) and absolute value of log_2_ fold change (FDR<0.01 and żlog2FCż > log(1.1,2)). SVM model from “e1071” package and “pROC” package were used to predict gender and plot ROC, respectively.

### Statistics

Data analysis was performed in R (version 3.6.0). Coefficient of variation (CV) of feature intensities in QCs was calculated and compared using t-test using a predefined significant level (*P-value<0.05*).

## Data availability

The raw data required to reproduce these findings cannot be shared at this time due to legal or ethical reasons.

## Code availability

R package and corresponding tutorial can be found at https://github.com/DLI-ShenZhen/RFQI. It is needed to be installed under linux environment.

## Author contributions

EL conceived the algorithm, wrote the code, analyzed data and wrote the draft of the manuscript. KC wrote the first draft of the manuscript and contributed to the study design. QL conducted data analysis, contributed to make graphs and developed R package. DL contributed to the method development of MS2 identification. BG conducted data analysis of gender classification. FL collected samples of ICXurineDB and FT conducted experiment of ICXurineDB. WL, YW and XY conducted experiments of MS2 validation. ZL, BL, and JS contributed to the study design and manuscript writing. HZ conceived the study, supervised the data analysis and contributed to the draft writing. All authors contributed to interpretation of the results. All authors read and approved the manuscript.

## Competing interests

iCarbonX has filed one patent on the basis of this study to the State Intellectual Property

Office of China **(2019102560903).**

The software is free for research use.

---

**Figure S1.**
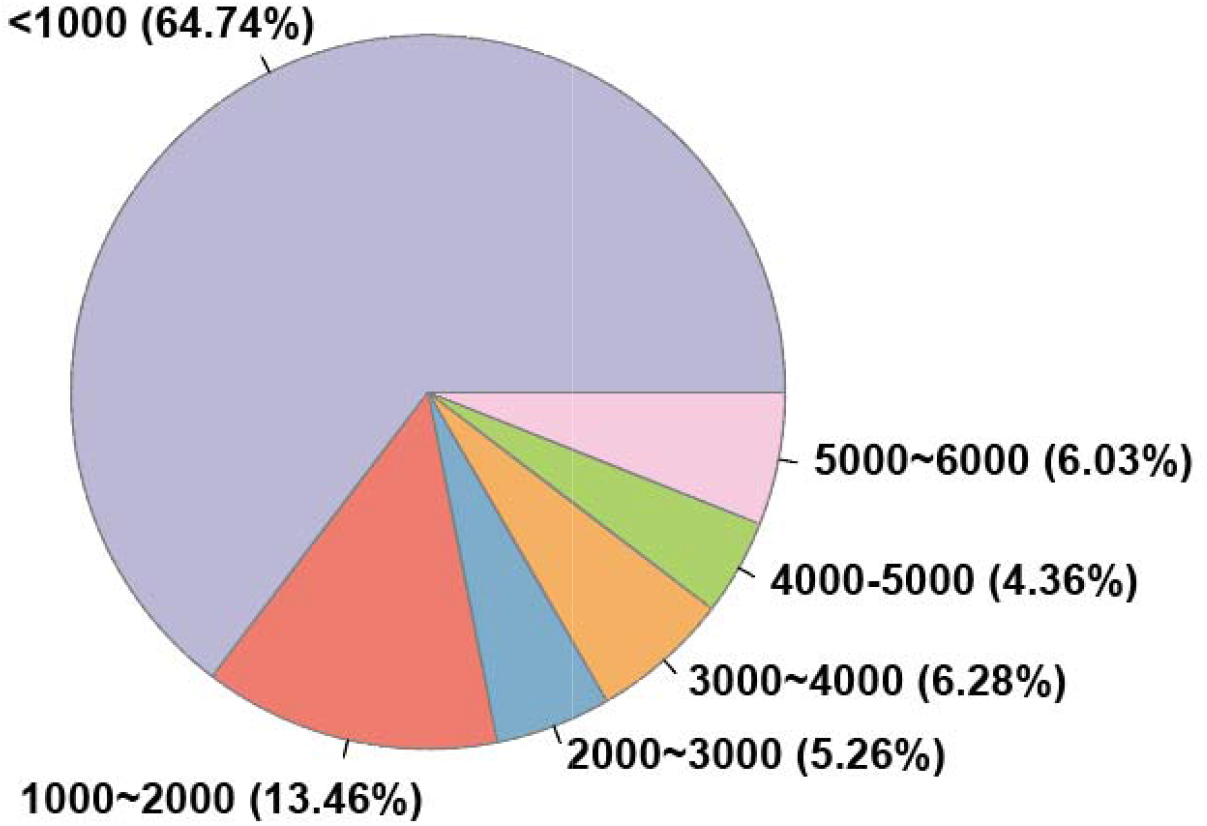
Metabolites inconsistency in 140 batches containing 6063 samples across 1.5 years. <1000 (64.74%) means that 64.74% of metabolites are detected in less than 1000 samples.

This plot is gif format, you need to view it in PPT play mode

**Figure S2.**
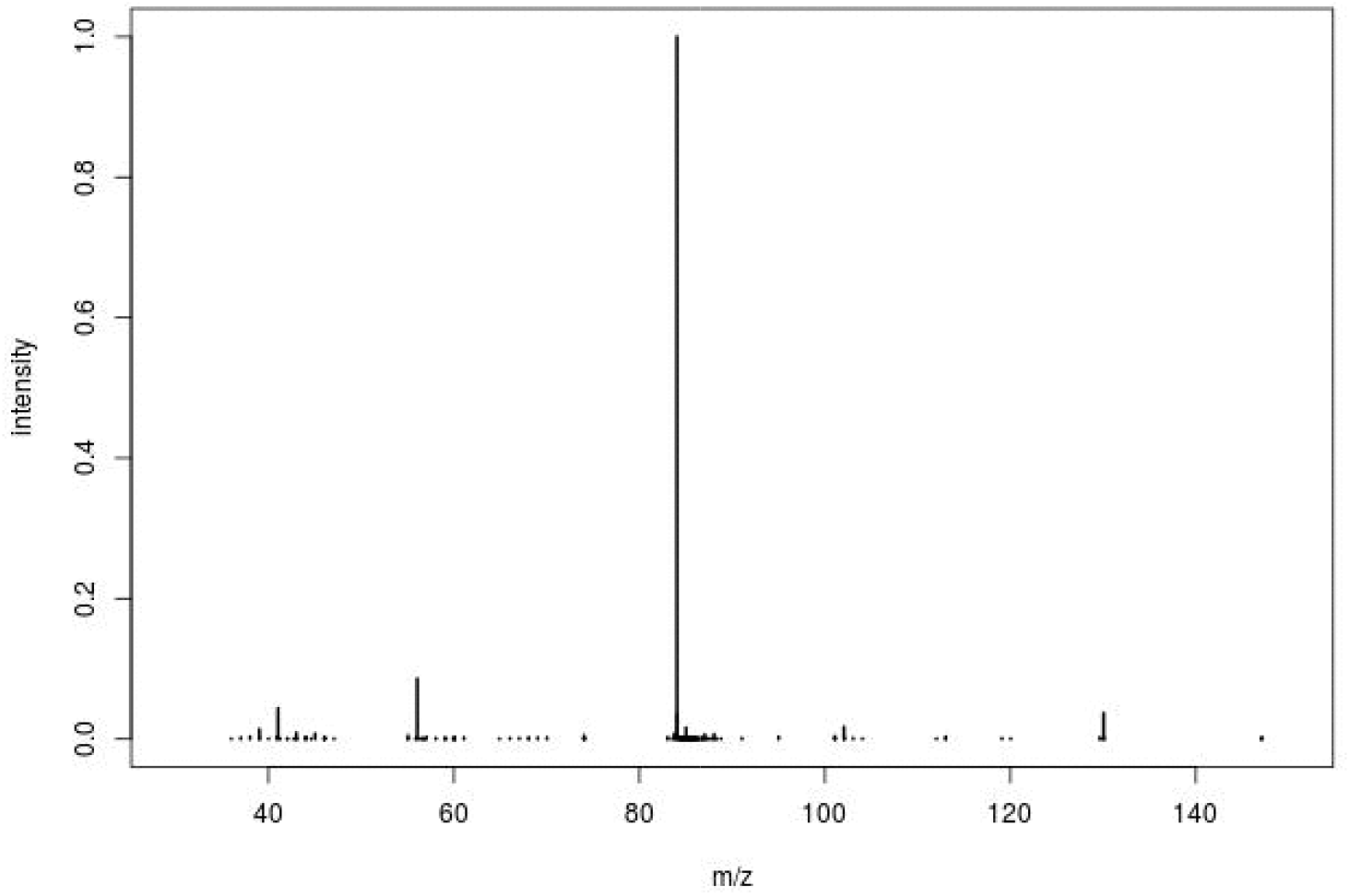
CID of a precursor is not stable. Intensities of m/z are fluctruate. (This plot is in gif format and should be viewed in PPT play mode)

**Figure S3.**
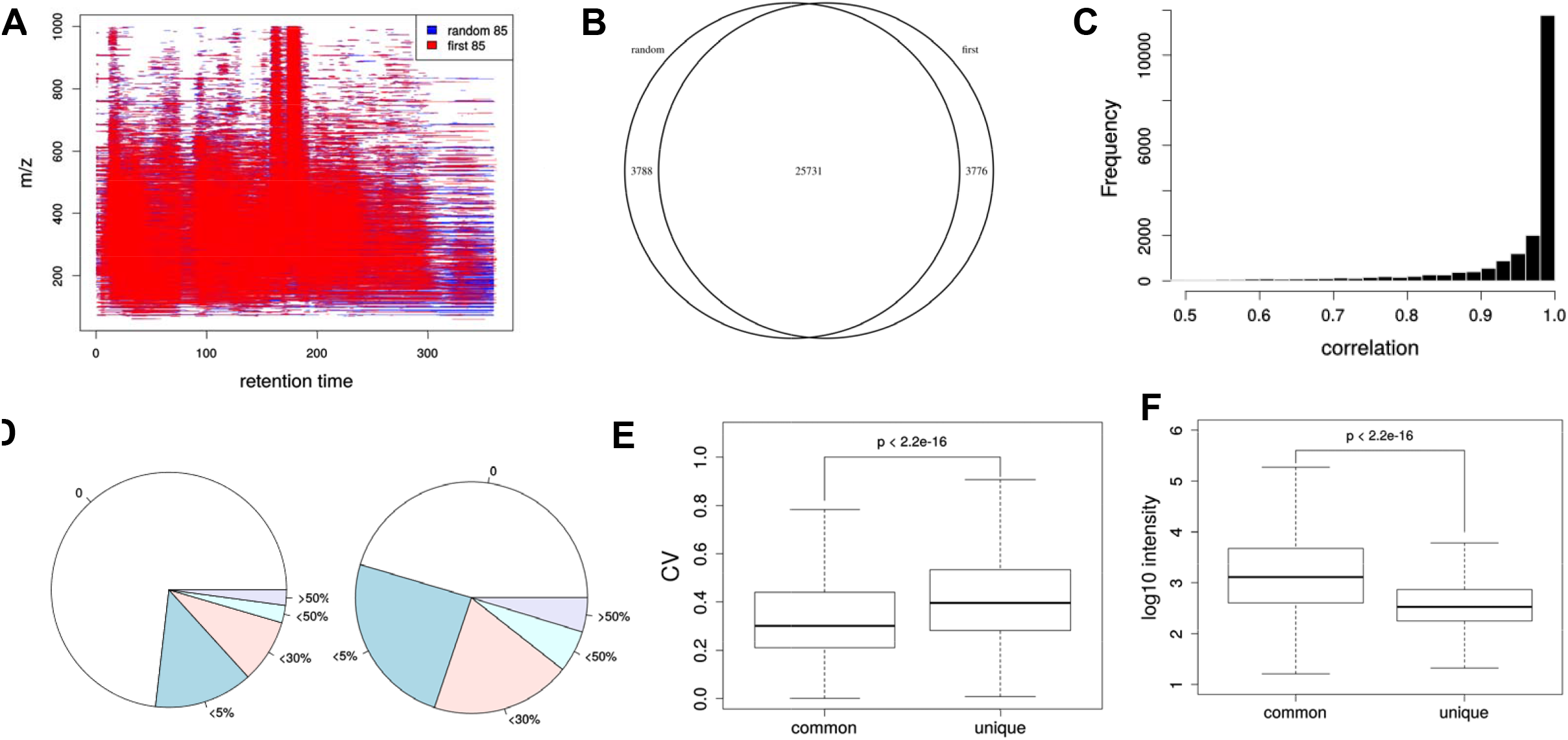
Comparison of serial rFT and random rFT. A. Feature area overlap. B. Featue overlap. C. Sample correlation of quantities based on one-to-one mapped features of serial rFT and random rFT. D. Missing value distribution of common features and unique featuers of serial rFT and random rFT. “0” represents features have 0 mssing value. E. CV comparison of common features and unique features. F. intensity comparison of common features and unique features.

**Figure S4.**
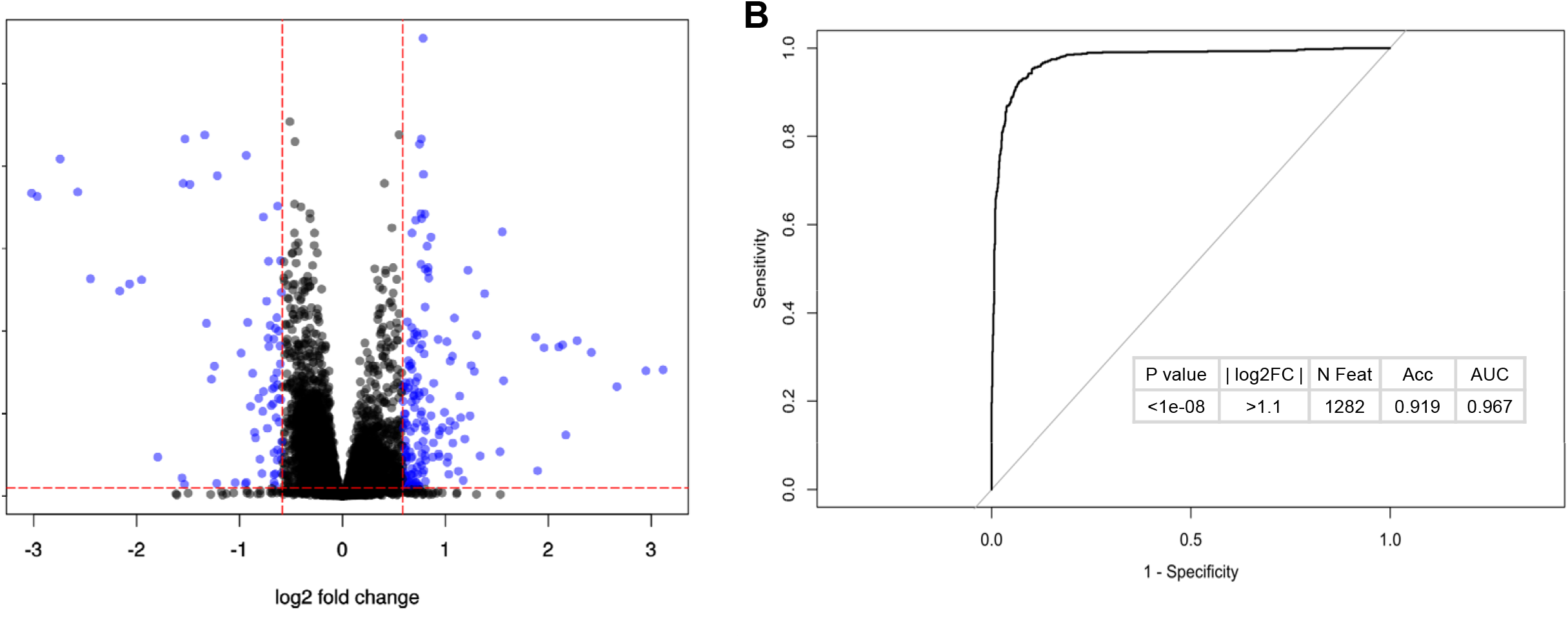
Classification of sex using real-time-analysis urine metabolic profile. A. Using volcano plot to select features. Only features with MS2 spectrum (10649) were used to do feature selection. Blue points were selected for classification. The horizonal dash red line represents fdr=0.01, the vertical dash red line represents | log2FC | = 1.5. B. ROC curve of classification of gender using support vector machine (svm) model.

